# Pooled CRISPR screening in pancreatic cancer cells implicates co-repressor complexes as a cause of multiple drug resistance via regulation of epithelial-to-mesenchymal transition

**DOI:** 10.1101/648709

**Authors:** Ryne C. Ramaker, Andrew A. Hardigan, Emily R. Gordon, Carter A. Wright, Richard M. Myers, Sara J. Cooper

**Author notes:** These authors have contributed equally to this work (Ryne C. Ramaker, Andrew A. Hardigan, Emily R. Gordon). Corresponding author: Sara J. Cooper.

## Abstract

Pancreatic ductal adenocarcinoma (PDAC) patients suffer poor outcomes in part due to therapeutic resistance. We conducted four genome-wide CRISPR activation (CRISPR_act_) and CRISPR knock out (CRISPR_ko_) screens to identify novel resistance mechanisms to four cytotoxic chemotherapies (gemcitabine, 5-fluorouracil, irinotecan, and oxaliplatin). ABCG2, a well-described efflux pump was the strongest mediator of resistance. We showed that overexpressing HDAC1 altered promoter occupancy and expression of genes involved in the epithelial-to-mesenchymal transition. Using the results of our CRISPR screens, we predicted drug sensitivity for patients and cell lines based on gene expression profiles. These predictions could be clinically useful for treatment selection.

## MAIN TEXT

Despite decades of work, pancreatic ductal adenocarcinoma (PDAC) has remained largely refractory to improvement of five-year survival rates, which are still less than 10% (1). Multi-drug combinations, such as FOLFIRINOX (fluorouracil, folinic acid, irinotecan and oxaliplatin), achieve, at best, modest improvements in patient outcomes (2). Unfortunately, many people with pancreatic cancer develop complete resistance to potent multi-drug cocktails (3). Cellular mechanisms of resistance have been explored by previous insertional mutagenesis- and RNA interference-based screens and have successfully identified genes whose inactivation leads to gemcitabine sensitivity in PDAC cells (4–6). Genome-wide CRISPR-Cas9 screening can provide complementary information (7, 8) and has identified essential genes in cell lines with *RNF43*-mutations (a recurrently mutated gene in PDAC) (9). These studies highlight the various mechanisms of cellular resistance and hint at the roles that genetic background and heterogeneity within tumors play in drug resistance (10).

The growing field of precision oncology aims to predict an optimal treatment for a patient based on tumor profiling. However, in the context of highly heterogeneous tumors, detection of genetic signatures associated with treatment response is difficult (11). One approach to this problem is to define the landscape of cellular mechanisms of PDAC drug resistance experimentally, then deeply screen tumors using a targeted approach for the presence of previously-identified resistance drivers. To achieve this goal, we performed CRISPR-Cas9 knock-out (CRISPR_ko_) (12) and endogenous activation (CRISPR_act_) (13) screening of 23,728 genes and 138,188 sgRNAs in two PDAC cell lines (BxPC3 and Panc-1) to identify genes whose loss or gain of expression were able to modulate sensitivity to four of the most common cytotoxic chemotherapies used in the treatment of PDAC (gemcitabine, oxaliplatin, irinotecan, and 5-fluorouracil, **Figure 1A, S1A-B**).

**Figure 1.**
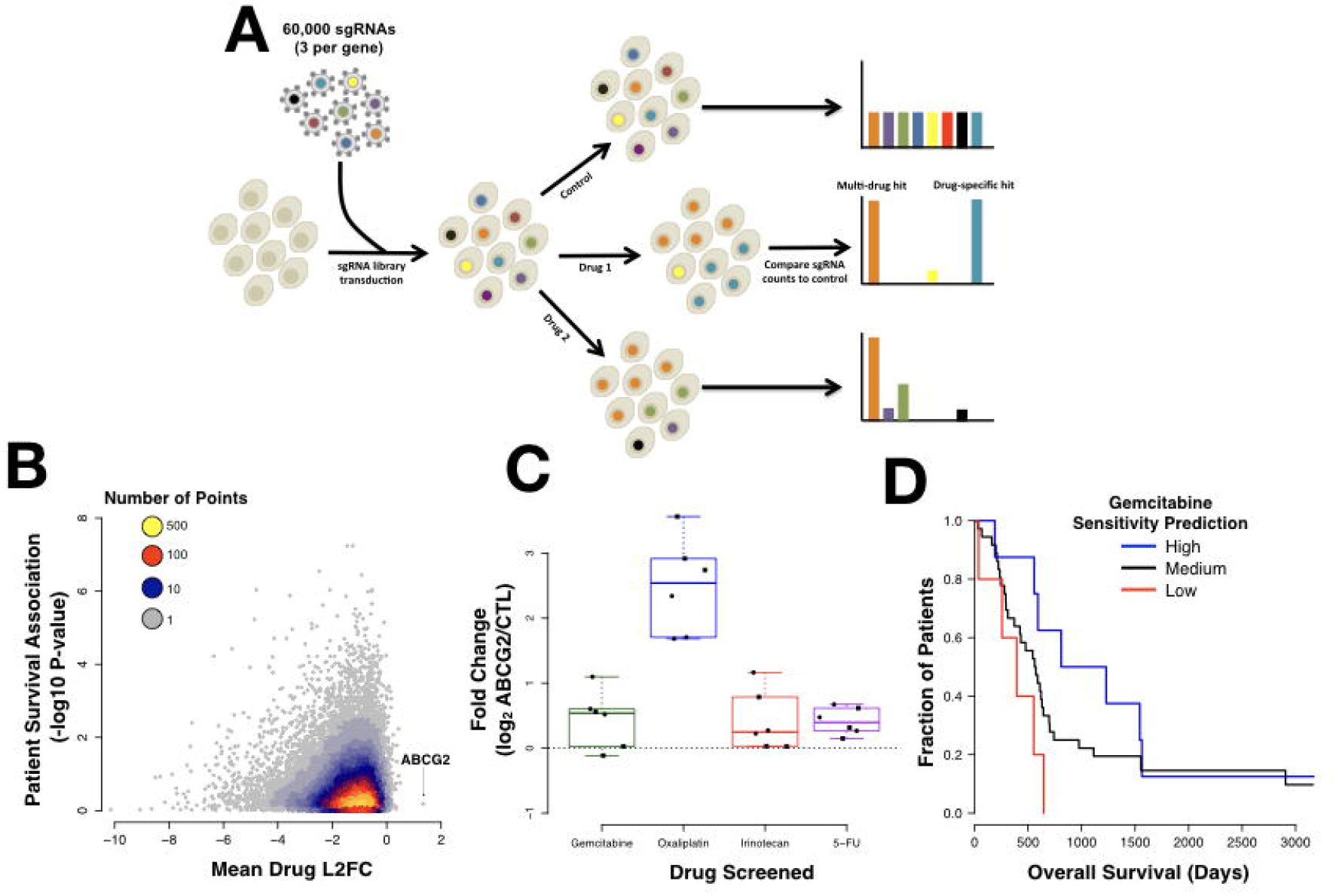
CRISPR screen reveals drug resistance genes. (A) Schematic describing our screening protocol. (B) A scatterplot shows the mean L2FC sum for all four drugs assayed in each of two cell lines compared to the log10 p-value for the association of the same gene’s expression with with patient survival. ABCG2 stands out as the most highest L2FC over all four drugs (C) Boxplots indicate the sgRNA fold change in counts per million comparing treated cells to control cells for each replicate and each cell line for the top *ABCG2* sgRNA. Circles represent data from Panc-1 cells. Squares represent BXPC-3 (D) Using weighted averages derived from the screen data we predicted likely sensitivity to gemcitabine based on expression of resistance-associated genes (PKG). There is a significant difference in survival between patients with predicted high versus low gemcitabine sensitivity (p=0.01, Chi-squared test).

We found the sgRNAs most strongly associated with drug resistance were highly drug-specific (Figure S1B) and replicates were significantly more correlated than samples treated with a different drug (**Figure S1C-F**). However, there was a much higher correlation between samples treated with different drugs than expected by chance, suggesting that mechanisms of resistance were sometimes shared between drugs in our study (**Figure S1C-F**). To prioritize resistance genes, we computed the sum of the replicate-minimum, log_2_ count fold change of the two most enriched sgRNAs targeting each gene (L2FC sum) in each cell line. We identified multi-drug resistance genes by computing the mean L2FC sum across all four drugs in each cell line (**Supplemental Table 1A-B**). This approach was particularly powerful because it leveraged information from 48 perturbations (4 drug screens x 3 replicates x 2 cell lines x 2 sgRNAs).

CRISPR_act_ of the ATP-binding cassette (ABC) transporter, ABCG2, was the only perturbation that persistently induced resistance across each of the drug treatments in both of our cell lines (**Figure 1B-C**). Individual follow up of our top ABCG2 sgRNA showed that it induced a highly specific, 30-fold overexpression of ABCG2 (**Figure S2A**) and produced a resistance phenotype that was reversed with previously developed inhibitors of ABCG2 (**Figure S2B-C**). ABCG2 functions as an efflux pump with a broad range of substrates and has been associated with multi-drug resistance in several previous studies (14–16); this provides a strong validation of the efficacy of our multi-drug screening approach. Despite being the strongest signal associated with multi-drug resistance in our screen, ABCG2 does not appear to have significant relevance in PDAC patient tumors, as it is expressed at relatively low levels in all prognosis groups (**Figure S2D**). However, further analysis suggests that our screen has identified additional patient-relevant resistance genes (**Figure 1B**). We computed drug sensitivity scores based on weighted expression level of resistance genes identified by our screen. Our algorithm PancDS is publicly available (https://github.com/rramaker/PancDS) and shows that those predictions readily segregate cell lines and patients into different treatment response groups (**Figure 1D, S3A-B**).

To identify other plausible mechanisms of multi-drug resistance with clinical relevance, we assessed whether there was enrichment of gene pathways associated with drug resistance or sensitivity based on our CRISPR_act_ and CRISPR_ko_ screens. Activation of chromatin remodeling genes was one of the most consistent mechanisms of drug resistance (**Figure 2A, Table S2**). Widespread chromatin repression has been previously associated with poor prognosis in PDAC patients (17). We performed targeted experiments with HDAC1, a gene that is known to cooperate with trans-acting repressors as a member of several transcriptional repressor complexes. Our top HDAC1 sgRNA induced greater than 10-fold overexpression of HDAC1. As one might expect, over-expression of this transcriptional regulator also produced several weaker transcriptional changes that we hypothesize to be downstream of HDAC1-mediated transcriptional regulation (**Figure 2B**). These downstream gene expression changes were particularly enriched for genes implicated in the epithelial-to-mesenchymal transition (EMT), a pathway known to mediate multi-drug resistance (**Figure 2B-C**) (18). Given HDAC1’s function as a member of canonical repressor complexes, we were surprised to find that its activation also resulted in several up-regulated genes, including IMP2, TIMP1, ANXA1, and WNK1, which are involved in promoting a mesenchymal or stem cell state (19–22). CRISPR_act_ of 11 other transcriptional repressors that were associated with resistance in our genome-wide screen (ARID4A, SMARCA4, SIN3A, SIN3B, SAP30, SAP18, RBBP7, MTA2, GATA2, CHD4, and BRMS1) also showed concordant up-regulation of at least one, but often several, of the same top genes over-expressed upon HDAC1-activation (**Figure 2D**). ChIP-sequencing (ChIP-seq) of cells with activated HDAC1 uncovered a 3-fold increase the number of occupied sites relative to control cells and identified 17,501 occupied sites specific to HDAC1 activation (**Figure 2E**). These HDAC1-specific ChIP-seq peaks were highly enriched near the transcription start sites of genes differentially expressed upon HDAC1 activation (**Figure 2F**, **Figure S4**), suggesting a role for direct binding of HDAC1 in modulating their expression. Supporting this observed HDAC1 gene regulatory network, we found HDAC1-activated cells exhibited more cell migration in scratch assays relative to control cells (**Figure 2G-H**).

**Figure 2.**
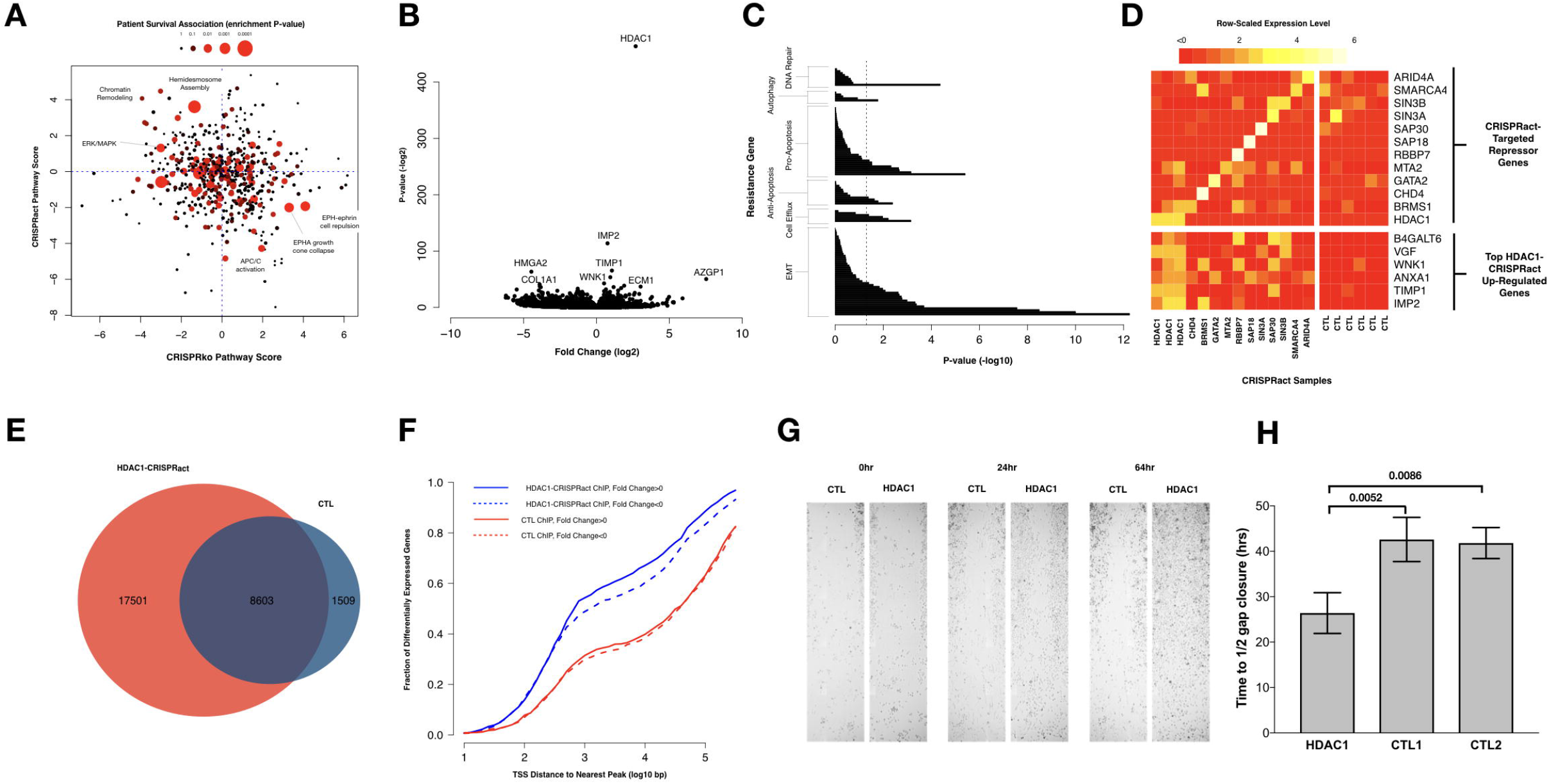
A) Scatter plot showing pathways enriched for multi-drug resistance in our CRISPR_act_ and CRISPR_ko_ screens and their association with patient survival in the TCGA cohort. Patient survival is represented by the size and color of the circle. B) A volcano plot shows that activation of HDAC1 expression using a dCas9-activation approach results in strong overexpression of HDAC1 based on RNA-sequencing. (C) Pathway enrichment analysis shows that HDAC1 overexpression especially affects epithelial to mesenchymal transition (EMT), cell efflux, apoptosis, autophagy, and DNA repair. P-values reported are derived from Fisher’s exact test comparing observed versus expected number of genes in each pathway. Full pathway list available in Table S2. (D) Overexpression of each of the target genes listed on top right quadrant leads to a similar pattern of over-expression of genes shown on the lower right. (E) ChIP-seq analysis reveals ChIP-seq peaks in the control (CTL) MiaPaCa-2 cells overlap significantly with MiaPaCa-2 cells over-expressing HDAC1 (red). An additional 17501 peaks are identified with over-expression of HDAC1. (F) Cumulative distribution plot showing that HDAC1 binding sites identified upon over-expression of HDAC1 (blue) are nearby transcription start sites (TSS) of differentially expressed genes. (G) Scratch assay shows that over-expression of HDAC1 leads to increased migration compared to control cells (H) Quantification of the scratch assays shows a significant difference with HDAC1 overexpression (Student’s T-test).

These data demonstrate the value of pooled CRISPR screening as a method for discovering cellular mechanisms of drug resistance. CRISPR_act_ of *ABCG2* was the highest-confidence perturbation capable of inducing multi-drug resistance, but its clinical relevance is unclear. Some reports indicate that *ABCG2* is expressed uniquely in cancer stem cells, a minority of the total cell population; thus, *ABCG2*’s role may be obscured in bulk tumor sequencing (16). Leveraging our screen data to predict drug sensitivity in cell lines and patients based on their respective gene expression profiles demonstrate the potential application of these data, en masse, to direct personalized therapeutic approaches.

Our pathway-based analysis linked CRISPR_act_ of several transcriptional repressor complex members to chemoresistance. Several chromatin remodeling genes, including *HDAC1*, induced a program of gene expression associated with EMT. The EMT pathway has been previously associated with chemoresistance in mouse models of PDAC where key EMT transcription factors were ablated (23). Chromatin remodeling genes have been linked to clinically relevant phenotypes in PDAC (24) including induction of EMT and increasing stem cell populations (25, 26). Our data provide evidence for *HDAC1* occupying the promoter regions and regulating expression of several genes over-expressed upon CRISPR_act_. Despite its canonical role as a repressor, HDAC complexes have recently been found to play a variety of roles in gene regulation (27). HDAC inhibitors have exhibited inconsistent results in clinical trials to date, an observation that has been partially attributed to a poor understanding of which HDAC classes are the best targets and which biomarkers indicate sensitivity to HDAC inhibition (28).

These data further our understanding of HDAC1 specifically and chromatin remodeling more generally in drug resistance and importantly the results of our large-scale screen provide additional avenues for exploration in the quest to improve treatment options for pancreatic cancer patients.

## METHODS

Panc-1 (CRL-1469), BxPC3 (CRL-1687), and MiaPaca-2 (CRL-1420) cell lines were obtained from ATCC and cultured according to ATCC specifications. LentiCas9-Blast (Addgene #52962) or Lenti-dCAS9-VPS46-Blast (Addgene #61425) with lenti-MS2-p65-HSF1-Hygro (Addgene #61426) were used to generated cells stably expressing the knockout or gene activation machinery, respectively. The GeCKO A pooled sgRNA library (Addgene #1000000049) was used for gene knock out screening and the SAM pooled sgRNA library (Addgene #1000000057) was used for gene activation screening. LentiSAMv2 (Addgene #75112) was used for single gene activation. sgRNA library cloning, viral packaging and transduction method was previously described (25). Primer sequences are available in Table S3.

Library-transduced cells were under selection for one week post-transduction and expanded to 7×10^7^ cells per treatment replicate or 1000x representation for each of the 4 drug (gemcitabine, oxaliplatin, irinotecan and 5-fluorouracil) and control conditions per replicate. A minimum 500x representation was maintained at all times in control cells. Drug treatment doses were optimized to yield ~80% cell death relative to untreated control cells after 14 days of culture. After 14 days of drug treatment, cells were pelleted and stored at −80°C. DNA extraction and library preparation was performed as previously described (29). Three sets of replicates (control and 4 drug treated samples) for each cell line were sequenced on one lane of Illumina NextSeq resulting in an average of 40 million reads per sample. We ranked each of the sgRNAs targeting each gene by the minimum log_2_ fold change across each replicate.

To identify top genes from our genome-wide screen, we prioritized genes by the “L2FC sum” in each cell line, which is the sum of the replicate minimum log_2_ fold changes of the top two sgRNAs targeting each gene. Multi-drug hits were prioritized by computing the mean “L2FC sum” of the four drug treatments.

Pathway enrichment significance was determined using a Wilcox-ranked sum test based on Reactome Pathways (**Table S2**) (30). Cell lines over-expressing chromatin remodelers, including HDAC1 were characterized using RNA-sequencing and ChIP-sequencing data generation and analysis using well established, published methods (31, 32) (https://www.encodeproject.org/documents/).

Cell migration was measured using scratch assays on MiaPaCa-2 cells. Time to close half the gap was calculated for each well (33).

Additional methodological details are available in the Supplemental Information.

## DATA AVAILABILITY

ABCG2 RNA-sequencing data are available at GEO using the accessions GSE131596.

RNA-sequencing and ChIP-sequencing data for chromatin remodelers are available using the GEO accession GSE158541.

Raw sequencing data from the screen are available through SRA using the project ID PRJNA542321.

## AUTHOR CONTRIBUTIONS

RCR, AAH, ERG, CAW and SJC designed the experiments. RCR, AAH, CAW, and ERG collected data. RCR, AAH, SJC, and ERG analyzed the data. RCR, AAH, and SJC wrote the first draft. All authors contributed to writing of the paper and read and approved the final manuscript.

## CONFLICT OF INTEREST DISCLOSURES

None reported.

## ACKNOWLEDGEMENTS

We thank the Myers and Cooper labs for helpful feedback. We also thank HudsonAlpha Discovery’s Genome Services Lab for assistance and optimizing the sequencing parameters of our CRIPSR screen. We also acknowledge The Cancer Genome Atlas, Cancer Cell Line Encyclopedia Project datasets, which were extremely valuable and without which this study would not be possible. This work was supported by the State Cancer Fund of Alabama (to RMM). RCR and AAH were funded by the UAB MSTP (NIH-NIGMS 5T32GM008361-21. SJC and ERG were supported by the HudsonAlpha Tie the Ribbons Fund. SJC received support from the UAB CCTS grant (NIH 1UL1TR001417-01) and the UAB Comprehensive Cancer Center (NIH 5P30CA013148).

## Notes

https://github.com/rramaker/PancDS

### Competing Interest Statement

The authors have declared no competing interest.

### Summary of Updates

Compared to the previous version, there are two main changes. First, we incorporate new ChIP-seq and RNA-sequencing data as well as functional characterization of HDAC1 over-expression in PDAC cell lines, providing further insight into the mechanisms of drug resistance and particularly how alterations in chromatin remodeling lead to dyregulation of the epithelial-to-mesenchymal transition, ultimately contributing to drug resistance. Second, we formalize using the results of the CRISPR screens to predict drug response based on gene expression data from patient tumor tissues and pancreatic cancer cell lines, highlighting the potential clinical applications for these data.

