## Supplementary figures and images for "Pooled CRISPR screening in pancreatic cancer cells implicates co-repressor complexes as a cause of multiple drug resistance via regulation of epithelial-to-mesenchymal transition"

### Supplemental Figure 1

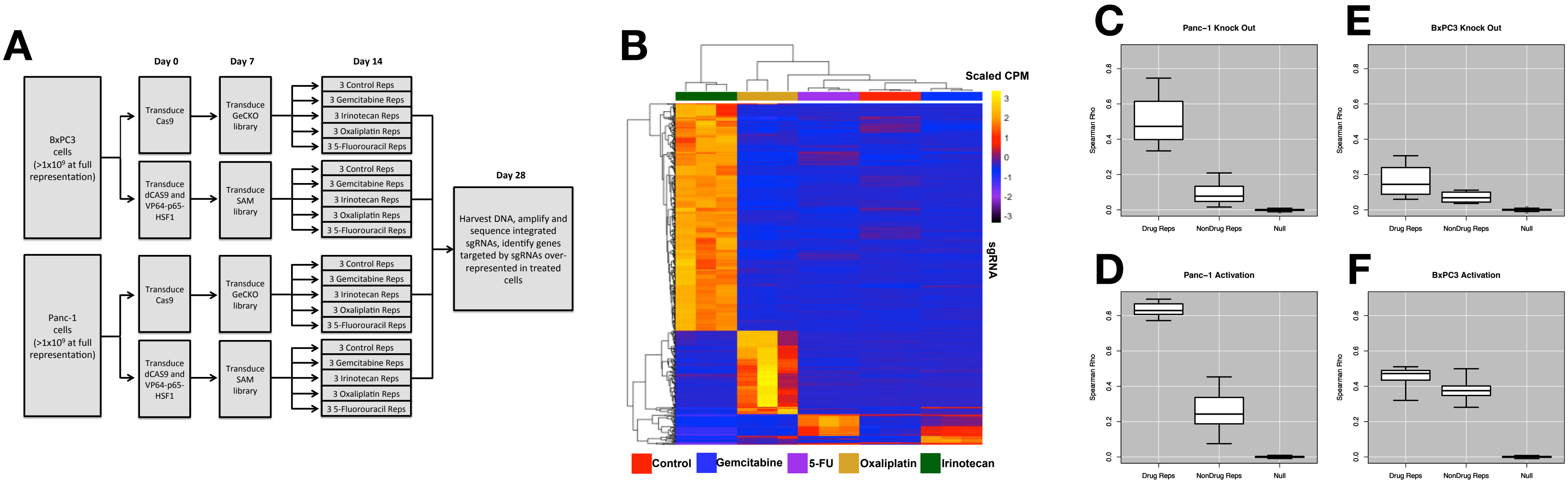

### Supplemental Figure 2

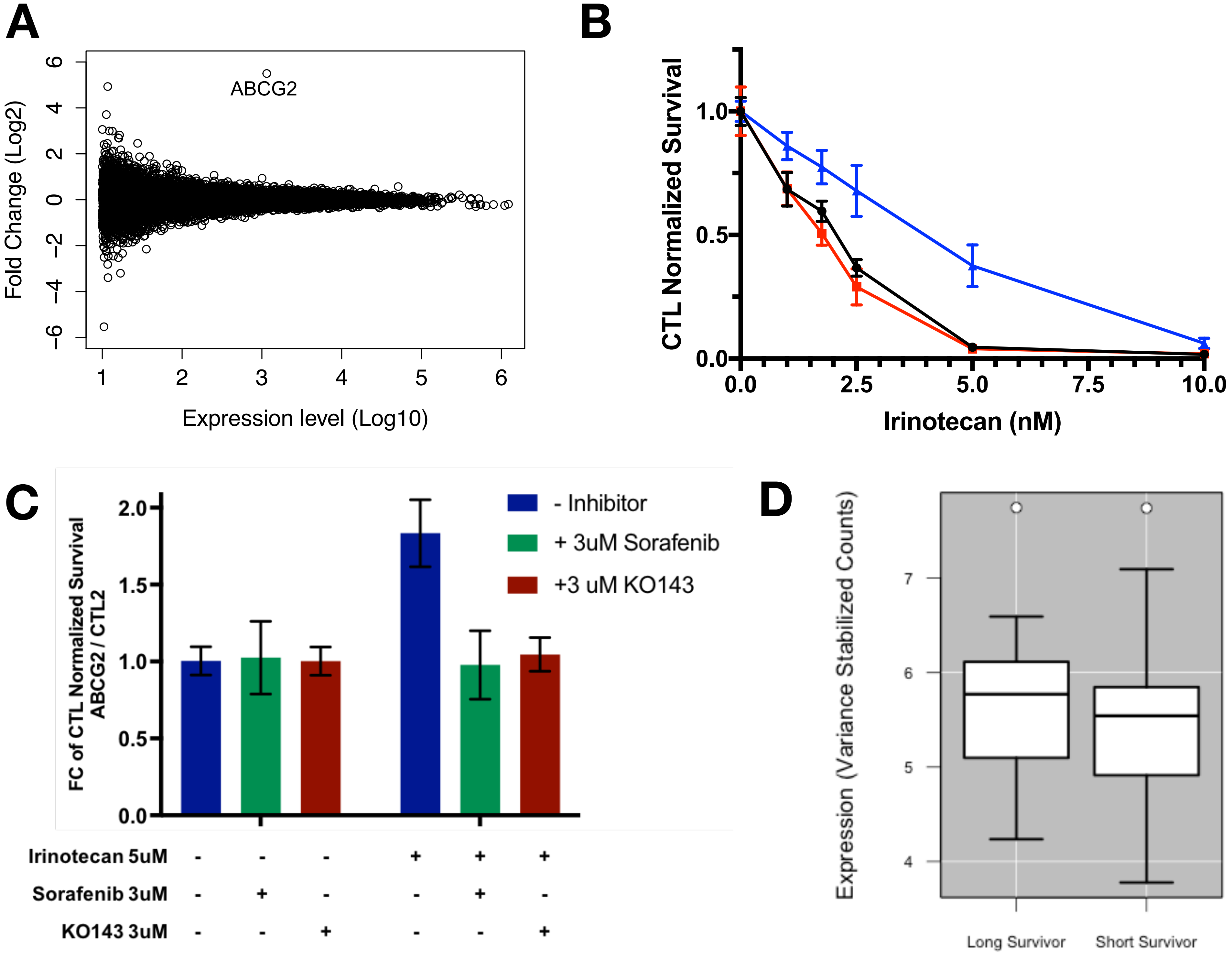

### Supplemental Figure 3

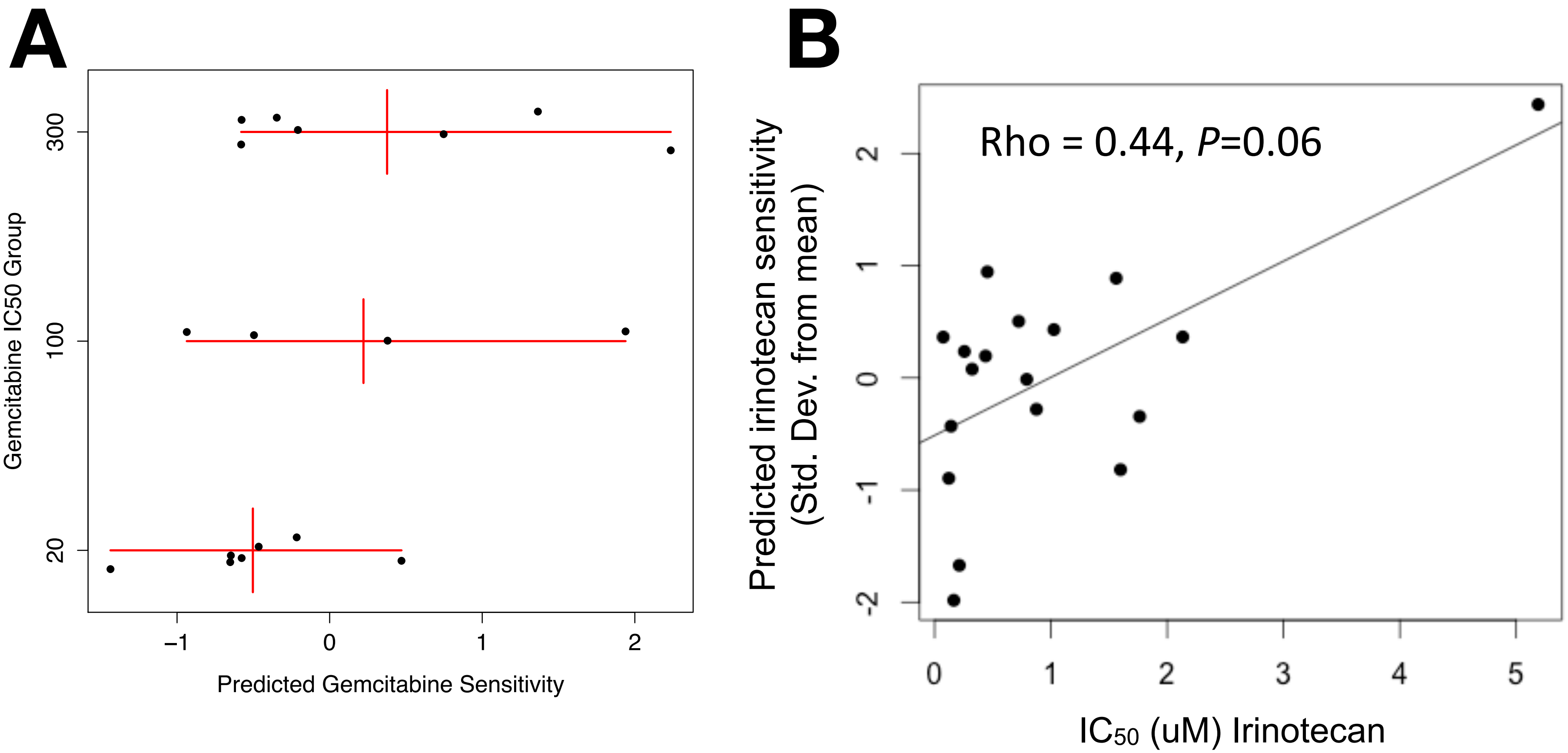

### Supplemental Figure 4

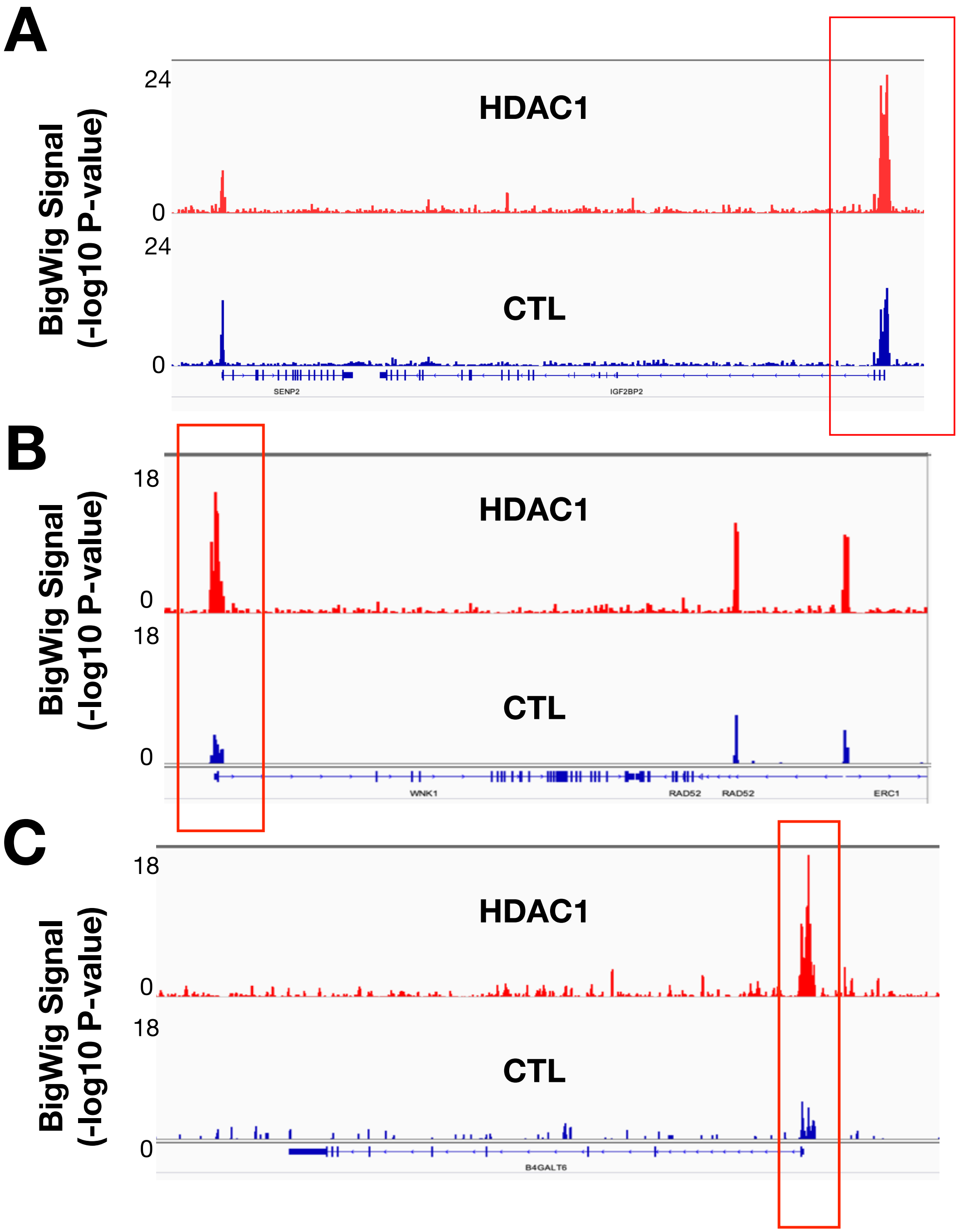
