## Supplemental_Information for "Pooled CRISPR screening in pancreatic cancer cells implicates co-repressor complexes as a cause of multiple drug resistance via regulation of epithelial-to-mesenchymal transition"

SUPPLEMENTAL METHODS

Key primer sequences for library construction are listed in Table S3.

Primer Sequences for individual gene over-expression are listed in Table S4

Screening conditions

Drug Concentrations for genome-wide screen

| | Panc-1 | BxPC3 |
|---|---|---|
| Gemcitabine Tocris #G6423 | 25 nM | 25 nM |
| Oxaliplatin Sigma #O9512 | 2.5uM | 2.5 uM |
| Irinotecan Sigma #I1406 | 500nM | 250nM |
| 5-Fluorouracil Sigma #F6627 | 7.5 uM | 5 uM |

ChIP-sequencing and RNA-sequencing analysis were completed on cells over-expressing HDAC1. ChIP-seq data were generated and analyzed as described (https://www.encodeproject.org/chip-seq/transcription_factor/) from $2x10^7$ cells with the two HDAC1 antibodies (Santa Cruz Biotech SC-81598 and Invitrogen PA1860). Libraries were sequenced on an Illumina NovaSeq instrument with an average of 33M 75bp single end reads.

Total RNA was extracted from 1M MiaPaca-2 and Panc-1 cells over-expressing the indicated genes (ABCG2, HDAC1, BRMS1, CHD4, GATA2, RBBP7,SAP18, SAP30, SIN3A, SIN3B, SMARCA4, ARID4A) using the Norgen Total RNA Purification Kit (Norgen #17200 and #25710). RNA quality was assessed using Qubit and Bioanalyzer (RIN range = 9.4-10). For ABCG2 over-expression lines, we used the NEB Next Ultra II polyA library prep kit (Cat# E7770) with 800ng of input. Samples were pooled and sequenced using an Illumina NovaSeq with paired end 100bp reads. For all samples over-expressing chromatin remodeling genes and relevant controls, 250ng of total RNA was used as input for Lexogen 3' mRNA-seq FWD library prep (cat #015).

We amplified libraries with 16 cycles of PCR. Libraries were pooled and sequenced on an Illumina NextSeq yielding an average of 4.6M single-end 75bp reads per sample.

Pathway analysis of the genome-wide screen was done to identify enriched gene sets among the top hits. Pathways enriched for genes conferring drug resistance were identified by comparing the distribution of $\log_2$ fold changes for the top three sgRNAs targeting each gene within a Reactome pathway to that of all other genes by Wilcox ranked sum test. Reactome pathways with fewer than 10 genes targeted in our libraries were excluded. A consolidated knock-out and activation score was computed for each pathway by summing the $-\log_{10}$ P-values for the top three sgRNAs of each pathway (**Table S2**).

ChIP-sequencing and data analysis were completed as previously published Briefly, HDAC1 over-expressing Mia-Paca-2 cells were cross-linked, harvested, and DNA was precipitated using two HDAC1 antibodies. Libraries were constructed, pooled and sequenced using Illumina NovaSeq single end 75bp reads. These data were generated and analyzed using published ENCODE protocols https://www.encodeproject.org/documents/.

RNA-sequencing and analysis for Panc-1 cells over-expressing ABCG2 was done as previously described (. RNA-sequencing libraries for MiaPaca-2 and Panc-1 cells were made used the Lexogen 3' RNA-sequencing kit (cat#). They were pooled and sequenced on Illumina NextSeq machine with 75bp single-end reads. Analysis used previously published methods including: trimgalore for adapter trimming, fastqc for quality score assessment, STAR for alignment to hg38, HTSeq-gene to map reads to features, and DESeq2 for differential expression analysis.

MiaPaca-2 cells were grown to perform scratch assays. Scratch assays were completed using standard growth conditions (ATCC) after plating $1x10^5$ cells. Cell migration was assessed using image captured every 8 hours on a Lionheart FX (Biomek). Based on 8 merged images collected over 64 hours, we calculated the time (hr) to close the gap by half the width as previously described (https://doi.org/10.1016/j.jid.2016.11.020). Images were compiled in R (version 3.5.0) and analyzed with GIMP (v. 2.10) and Image J (version 1.5.2q).

SUPPLEMENTAL FIGURE LEGENDS

**Figure S1**. (A) Schematic detailing the screening workflow. (B) A heatmap of the sgRNA count per million, z-scored by row for the most variable 500 sgRNAs across all of our CRISPRa screen replicates in the PANC-1 cell line. (C-F) Boxplots describing the replicate, non-replicate, and scrambled null permutation Spearman correlations for knockout screens in Panc-1 (C) and BxPC3 (E) and activation screens in Panc-1 (D) and BxPC3 (F).

**Figure S2**. (A) Scatterplot shows ABCG2 stands out as the single gene showing a highly significant change in expression with use of the ABCG2_A and ABCG2_B sgRNAs to activate ABCG2 expression. These data are generated from RNA-sequencing of MiaPaca-2 and Panc-1 cells over-expressing ABCG2. (B) Cell survival curves display the fraction of cells surviving after a series of irinotecan doses for MiaPaca2 cells stably expressing an *ABCG2*-targeting sgRNA (blue) or non-targeting control sgRNAs (red and black). C) Each bar represents a ratio of ABCG2 over-expressing cells compared to a non-targeting control with the indicated drug treatments (+ indicates a treatment was added). Treatment of MiaPaca2 cells over-expressing ABCG2 using the ABCG2 guide shows no significant cell death with inhibitor only, where as ABCG2 alone confers drug resistance. The combination of 3uM sorafenib (green) or KO143 (maroon) with ABCG2 over-expression restores normal sensitivity to Irinotecan. (D) Boxplots showing the normalized expression levels of *ABCG2* in Kirby et. al. for patients surviving <300 days or >900 days. P-value is not significant.

**Figure S3**. (A) Predicted Gemcitabine Sensitivity was calculated using a weighted average of gene expression for resistance-associated genes based on expression profiles for 18 pancreatic cancer cell lines. These data were plotted compared to experimentally measured IC50 groups. The Wilcoxon P-value between the most resistant group of cell lines and the most sensitive is 0.097. (B) Scatterplot showing observed irinotecan $IC_{50}$ values compared to predicted sensitivity for 18 PDAC cells assayed by the Cancer Cell Line Encyclopedia. (Rho=0.44, p=0.06).

**Figure S4** ChIP-seq analysis of HDAC1-overexpressing (red) MiaPaca-2 cells compared to non-targeting controls (blue). HDAC1 overexpressing cells show increased HDAC1 peak height at (A) the IGF2BP2 promoter, (B) the WNK1 promoter, and (C) the B4GALT6 promoter.

SUPPLEMENTAL TABLES

Table S1. L2FC sums for each sgRNA targeting each gene for each drug and cell line. A) L2FC sums for $CRISPR_{act}$ B) L2FC sums for $CRISPR_{ko}$

Table S2. Pathway enrichment P-values $CRISPR_{act}$ and $CRISPR_{ko}$ screens indicating enrichment of hits among each reactome pathway. TCGA P-value indicates whether genes from that pathway are significantly associated with patient survival.

Table S3. Primers used in library amplification.
